# A mechanism-based computational model to capture the interconnections among epithelial-mesenchymal transition, cancer stem cells and Notch-Jagged signaling

**DOI:** 10.1101/314187

**Authors:** Federico Bocci, Mohit K. Jolly, Jason George, Herbert Levine, José N. Onuchic

## Abstract

Epithelial-mesenchymal transition (EMT) and cancer stem cell formation (CSCs) are two fundamental and well-studied processes contributing to cancer metastasis and tumor relapse. Cells can undergo a partial EMT to attain a hybrid epithelial/mesenchymal (E/M) phenotype or a complete EMT to attain a mesenchymal one. Similarly, cells can reversibly gain or lose ‘stemness’. This plasticity in cell states is modulated by signaling pathways such as Notch. However, the interconnections among the cell states enabled by EMT, CSCs and Notch signaling remain elusive. Here, we devise a computational model to investigate the coupling among the core decision-making circuits for EMT, CSCs and the Notch pathway. Our model predicts that hybrid E/M cells are most likely to associate with stemness traits and exhibit enhanced Notch-Jagged signaling – a pathway that is implicated in therapeutic resistance. Further, we show that the position of the ‘stemness window’ on the ‘EMT axis’ is varied by altering the coupling strength between EMT and CSC circuits, and/or modulating Notch signaling. Finally, we analyze the gene expression profile of CSCs from several cancer types and observe a heterogeneous distribution along the ‘EMT axis’, suggesting that different subsets of CSCs may exist with varying phenotypes along the epithelial-mesenchymal plasticity axis. Our computational model offers insights into the complex EMT-stemness interplay and provides a platform to investigate the effects of therapeutic perturbations such as treatment with metformin, a common anti-diabetic drug that has been associated with decreased cancer incidence and increased lifespan of patients. Our mechanism-based model helps explain how metformin can both inhibit EMT and blunt the aggressive potential of CSCs simultaneously, by driving the cells out of a hybrid E/M stem-like state with enhanced Notch-Jagged signaling.

## Introduction

Metastatic spread of cancer cells claims the highest number of fatalities, accounting for over 90% of cancer-related deaths[1]. Studies in mouse models have suggested that a large percentage of metastases are formed by clusters of Circulating Tumor Cells (CTCs) – cohesive units of more than two CTCs that are launched into the bloodstream as aggregates [2]. Consistently, clinical data highlights that the presence of clusters of CTCs correlate with higher aggressiveness and shorter patient survival across cancer types [3]. Thus, understanding the mechanisms that contribute to the formation and enhanced metastatic ability of these clusters holds promise for unraveling novel therapeutic strategies.

To enter the bloodstream as clusters of CTCs, epithelial cancer cells in primary solid tumors typically partially lose their cell-cell adhesion with their neighbors, and simultaneously acquire mesenchymal traits of motility and invasion. Such a hybrid epithelial/mesenchymal (E/M) phenotype, also referred to as a partial epithelial-mesenchymal transition (pEMT) state, facilitates clustered or collective cell migration [4,5]. A core regulatory circuit that receives multiple inputs and controls many molecular and morphological aspects of EMT consists of two families of microRNAs (miR-34 and miR-200) and two families of EMT-inducing transcription factors (EMT-TFs) (SNAIL and ZEB) [6] (Fig. 1, EMT module). High levels of miR-34 and miR-200, and low levels of SNAIL and ZEB correspond to an epithelial (E) state; an opposite configuration with (low miR-34 and miR-200, high SNAIL and ZEB) correspond to a mesenchymal (M) state [7–9]. An intermediate expression of these microRNAs and EMT-TFs has been proposed to correspond to a hybrid E/M state, exhibiting both cell-cell adhesion and cell motility [4].

**Figure 1.**
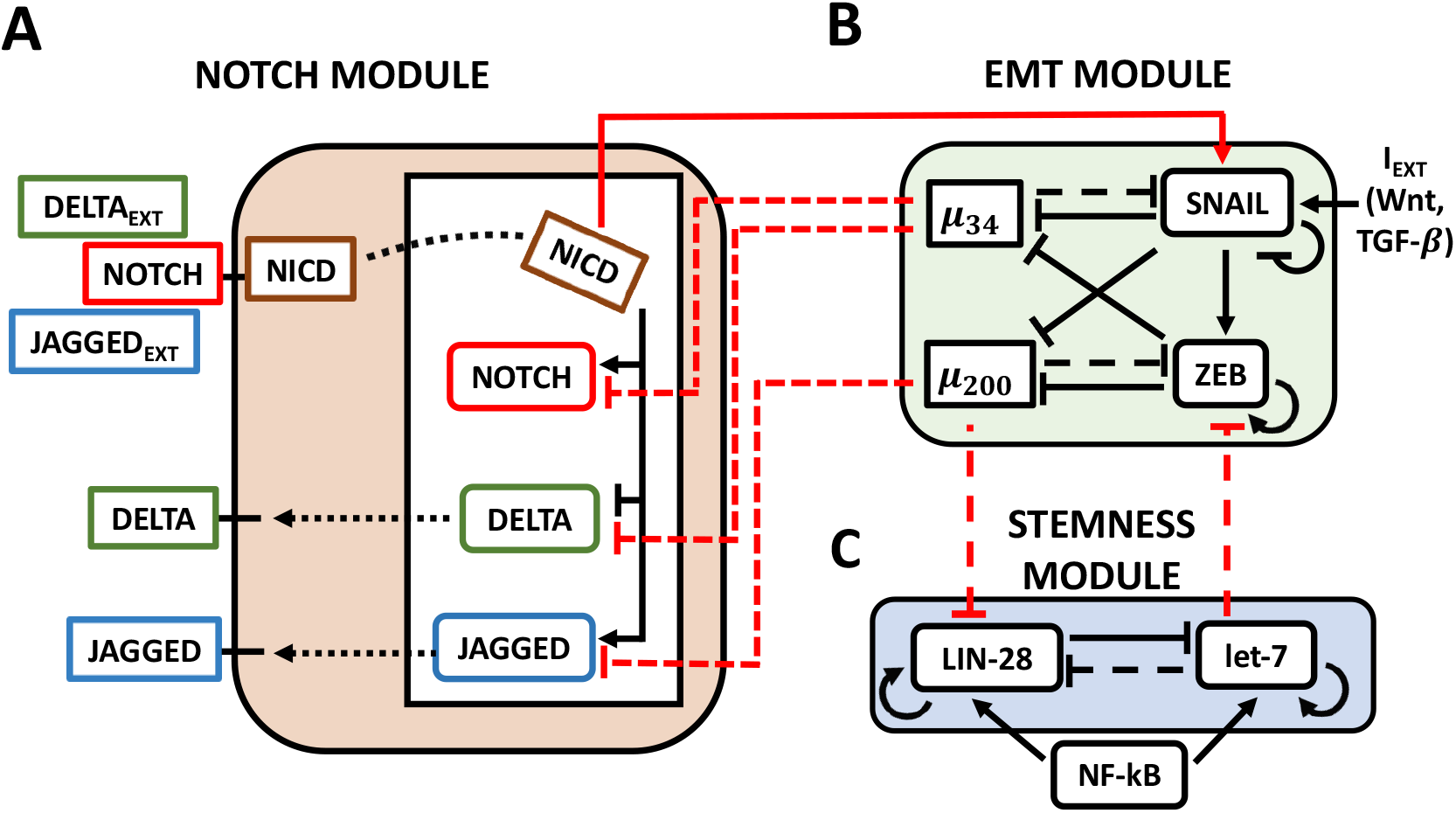
The Notch-EMT-STEM decision-making circuit. **(A)** The Notch circuit receives external ligands (Delta/Jagged) as inputs that bind to Notch on the surface. This results in the cleavage of Notch which generates NICD. NICD translocates to the nucleus where it transcriptionally activates Notch and Jagged while inhibiting Delta. **(B)** The EMT module contains two micro-RNA families (miR-34, miR-200) and two transcription factor families (SNAIL, ZEB) which mutually repress each other. External signals such as Wnt or TGF-beta activate SNAIL and promote EMT. **(C)** In the STEM module, LIN-28 and let-7 mutually repress each other, while both can self-activate. Additionally, external NF-kB signaling activates both. Solid lines stand for transcriptional/translational interactions, while dotted lines represent post-translational inhibition. The connections between the modules are highlighted in red: NICD transcriptionally activates SNAIL; miR-34 post-translationally inhibits Notch and Delta, while miR-200 inhibits Jagged; miR-200 and let-7 post-translationally inhibit LIN-28 and ZEB respectively.

Once individual CTCs and/or CTC clusters exit the bloodstream at a distant organ, they need to form secondary tumors. All three phenotypes – epithelial, mesenchymal, and hybrid E/M - have been correlated with possessing stemness, i.e. tumor-initiation ability [10–12], in different systems. Thus, the ‘stemness window’ can move along the EMT axis [13], and a precise mechanistic connection between EMT and cancer stem cells (CSCs) remains elusive. In many cases, a mutually inhibitory feedback loop between LIN-28 and let-7 regulates the tendency of a cell to behave as a CSC [14]. This loop can behave as a three-way switch [15] by giving rise to three possible states: (i) a (low LIN-28, high let-7), or DOWN (D) state; (ii) a (high LIN-28, low let-7), or UP (U) state; and (iii) a (medium LIN-28, medium let-7), or DOWN/UP (D/U) state (Fig. 1, STEM module). Since intermediate levels of OCT4, a direct target of LIN-28, has been observed to correlate with stem-like properties [16,17], the D/U state was proposed to be associated with stemness.

Additionally, cell-cell signaling through the Notch signaling pathway has been implicated in modulating EMT, enhancing therapeutic resistance[18], expanding the CSC population[19–21], and in the formation of clusters of CTCs [22,23]. Notch signaling is an evolutionarily conserved cell-cell signaling pathway[24] which includes a family of trans-membrane receptors (Notch) and two families of ligands (Delta, Jagged). The binding of one cell’s receptor to a neighbor cell’s ligand results in the cleavage of the Notch Intra-Cellular Domain (NICD), which translocates to the nucleus and regulates several target genes, including activating Notch and Jagged, but repressing Delta (Fig. 1, Notch module). In the presence of strong Notch-Delta signaling, a cell can either attain a (high Notch, low Delta) Receiver (R) state or a (low Notch, High Delta) Sender (S) state. Conversely, predominance of signaling toward the Notch-Jagged pathway culminates in a (high Notch, high Jagged) Sender/Receiver (S/R) state[25,26].

Here, we develop a mechanism-based mathematical model to elucidate the interconnections between EMT, CSCs and Notch signaling, by investigating the emergent dynamics due to the coupling among the EMT, STEM and NOTCH modules. We find that stemness traits tend to co-exist with a hybrid epithelial-mesenchymal (E/M) phenotype and strong Notch-Jagged signaling. Modulation by external signaling pathways can decouple the abovementioned correlation, suggesting that stem-like traits need not be exclusively correlated with a specific EMT phenotype. To validate this prediction, we examine the gene expression profile of CSCs from several cancer subtypes and find a heterogeneous distribution for the ‘stemness window’ along the ‘EMT axis’, thus enabling the existence of subsets of CSCs with epithelial, hybrid E/M or mesenchymal phenotypes. Lastly, we apply our formalism to model the action of metformin in targeting CSCs and inhibiting EMT, providing a mechanism-based explanation for several experimental findings, including decreased Notch1 levels in metformin-treated pancreatic cancer cells[27], metformin inhibition of TGF-beta induced EMT[28] and the recovery of stem-like traits upon NF-kB overexpression in metformin-treated cells[29].

## Results

### A mathematical framework to couple Notch signaling, Epithelial-Mesenchymal Transition and Stemness

To elucidate the outcome of interconnections among Notch signaling, EMT and stemness, we hereby develop a mechanism-based mathematical framework that integrates the experimentally identified connections among these three crucial pro-metastatic modules, and explore any correspondence between the cell states enabled independently by these three modules.

Given that each of the modules – Notch, EMT and STEM – can give rise to three different cell states independently, their coupling can possibly give rise to as much as 27 = 3^3^ states combining the (S, S/R, R), (E, E/M, M) and (D, D/U, U) states. Coupling between the modules can, however, introduce correlations which lower this to a much small number. The three modules are connected as follows (see red arrows on Fig. 1): (a) Active Notch signaling (NICD) promotes EMT through activating EMT-TF SNAIL and therefore increasing its cellular production rate constant by a fold-change factor λ_SNAIL_ > 1 [30,31]; (b) miR-34 and miR-200 act as post-translational inhibitors of Notch receptor and ligands, therefore increasing their degradation rate, hence shunting the activation of the Notch pathway [32–34]; and (c) LIN28 and ZEB are post-translational targets of micro-RNAs miR-34 and let-7, respectively [13], thus, miR-34 and let-7 decrease the production rate constants of LIN-28 and ZEB by factors λ_LIM–28_, λ_ZEB_ < 1, respectively (see Methods).

To investigate the cell-fate dynamics of this coupled network, we set up a mathematical model for the EMT-STEM-Notch coupled circuit (see Methods). The output of our model is a cell phenotype defined as a combination of (EMT, STEM, Notch) phenotype, depending on baseline conditions and strengths of interactions among these three modules.

### Notch-Induced EMT couples a hybrid E/M phenotype with Stem-like properties and a Sender/Receiver state

As the first step to investigate the coupling among states of Notch signaling, EMT, and stemness (i.e. STEM module), we examined how an epithelial cell responds to varying levels of external Notch ligands *L_EXT_* (either Delta or Jagged).

Following the binding of Delta or Jagged to Notch, Notch signaling is activated and stimulates EMT, thereby decreasing the levels of miR-34 and miR-200, while increasing those of SNAIL and ZEB (Fig. 2A and S1). This progression to EMT is achieved in two steps – transition from an epithelial state to a hybrid E/M state, and transition from a hybrid E/M state to a mesenchymal state. Specifically, an intermediate exposure to Notch ligands may enable cells to stably maintain a hybrid E/M state (yellow shaded horizontal region in Fig. 2A). LIN28 is inhibited by miR-200 (Fig. 1) and is therefore upregulated by the external Notch stimulus (Fig. 2B). Interestingly, projecting the stability region of the hybrid E/M state onto the LIN28 bifurcation diagram revealed a significant overlap between a hybrid E/M state (yellow shaded vertical region in Fig. 2B) and intermediate LIN28 levels (violet shaded horizontal region in Fig. 2B) corresponding to a DOWN/UP (D/U), or stem-like, state. Further, this stem-like hybrid E/M phenotype overlaps significantly with high Notch Intracellular Domain (NICD), which in turn biases the cell toward a (high Notch, high Jagged), i.e. hybrid S/R phenotype (Fig. 2C).

**Figure 2.**
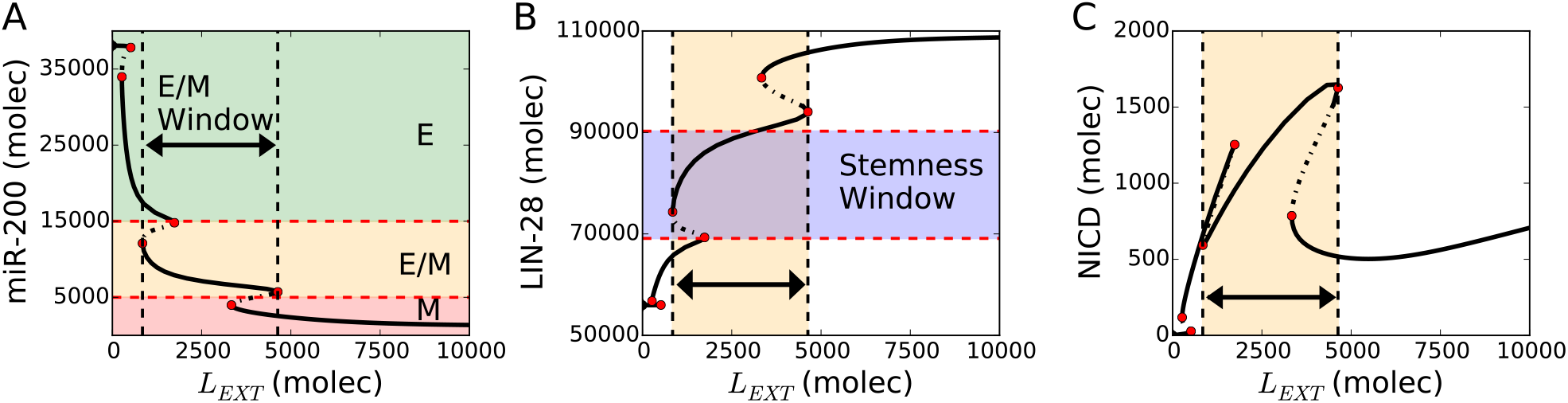
Notch-induced EMT couples hybrid E/M, stem and sender/receiver (S/R) states. **(A)** Bifurcation diagram of the cellular level of microRNA 200 (miR-200) as a function of external Notch ligands *L_EXT_* (Delta + Jagged). Continuous and dotted black lines denote stable and unstable solutions of the model respectively. The thick black horizontal arrow highlights the range of *L_EXT_* allowing a hybrid E/M state. Red dotted lines indicate the miR-200 range of epithelial, hybrid E/M and mesenchymal phenotypes. **(B)** Bifurcation diagram of the cellular levels of LIN-28 in response to *L_EXT_*. The cell falls into the stemness window at intermediate LIN-28 levels (violet-shaded horizontal region – see Methods for more details about defining the boundaries of stemness window). The yellow vertical region and black arrow highlight the range of *L_EXT_* levels enabling a stable hybrid E/M state. **(C)** Bifurcation diagram of NICD in response to *L_EXT_*. The hybrid E/M phenotype interval maps onto high levels of NICD corresponding to (high Notch, high Jagged) and therefore to a Sender/Receiver (S/R) phenotype (see Methods for more details about defining the boundaries of the Notch states). For this diagram, the coupling between EMT and STEM circuits is intermediate (*λ*_*LIM*–28_= *λ_ZEB_* = 0.5), external Notch receptor is *N_EXT_* = 10^4^ molecules and *NF* −*kB* = 2.5 10^3^ molecules.

Taken together, these results highlight a strong correlation between a hybrid S/R Notch state, a hybrid E/M phenotype, and the expression of stem-like traits, or a D/U state. In other words, cells in a hybrid E/M phenotype are highly likely to exhibit stem-like properties and show enhanced Notch-Jagged signaling. Indeed, cells co-expressing various epithelial and mesenchymal genes display enhanced JAG1 levels in circulating tumor cell (CTC) clusters and in drug-tolerant breast cancer cells [22]. Thus, this proposed overlap of hybrid E/M, stem-like traits, and Notch-Jagged signaling is supported by preliminary experimental evidence.

In the following sections, we will refer to the modes of the EMT module (E, E/M, M) as states because they have a one to one correspondence with the mathematical solutions of the model. In other words, each branch of the bifurcation diagram of Fig. 2A can be associated with a distinct EMT phenotype. Conversely, the modes of the Notch module (S, S/R, R) and STEM module (D, D/U, U) will be referred to as ‘phenotypes’ because they are defined based on threshold levels of (Notch, Jagged) and LIN-28, respectively rather than on the branches of the diagrams of Fig. 2B-C (see Methods for details).

### Varying the coupling strengths of EMT and STEM modules can enable shifts in the positioning of the “stemness window” on the EMT axis

To investigate the robustness of overlap among hybrid E/M, stem-like traits, and enhanced Notch-Jagged signaling, we vary the strengths of two links in our network: the cellular production fold-change of LIN-28 due to inhibition by miR-200 (*λ*_*LIM*−28_), and the cellular production fold-change of ZEB due to inhibition by let-7 (*λ_ZEB_*) [13]. These parameters can be varied from 0 (very strong repression) to 1 (no repression), generating a full spectrum, or diagram, of EMT-STEM coupling interactions. It should be noted that in our circuit, NOTCH and STEM modules are not directly coupled (Fig. 1).

First, we consider a cell exposed to an intermediate external Notch ligands. Such a cell will have intermediate NICD levels, and thus can attain a hybrid E/M state in the absence of connection between the EMT and STEM modules (Fig. 3A). The EMT and STEM diagrams highlight large parameter regions where the hybrid E/M and the D/U stem-like states are available to the cell, either as the only solution or as one of the two bistable solutions ({E/M}, {E, E/M} phases in Fig. 3B, and {D/U}, {D, D/U}, {D/U, U} phases in Fig. 3C). In particular, a strong repression of ZEB by let-7 pushes the cell toward an epithelial state (*λ_ZEB_* close to 0 in Fig. 3B and Fig. 3D), consistent with experimental observations that depleting ZEB1 can push the cells towards an epithelial phenotype[35]. Similarly, increasing the strength of repression of LIN28 by miR-200 (i.e. varying *λ_LIN−2_* from 1 to 0) induces a shift from a U to a D/U to a D state (Fig. 3C and Fig. 3D). Overlapping the EMT and STEM maps plotted above highlights a large {E/M – D/U} region (darker area in Fig. 3D)., i.e. a hybrid E/M state overlaps with a stem-like behavior. Additionally, a strong repression of ZEB by let-7 enables maintaining stem-like traits in an epithelial state {E – D/U}, while hybrid E/M states that are not stem-like can be observed for very strong {E/M – D} or very weak {E/M – U} inhibition of LIN-28 by miR-200 (Fig. 3D).

**Figure 3.**
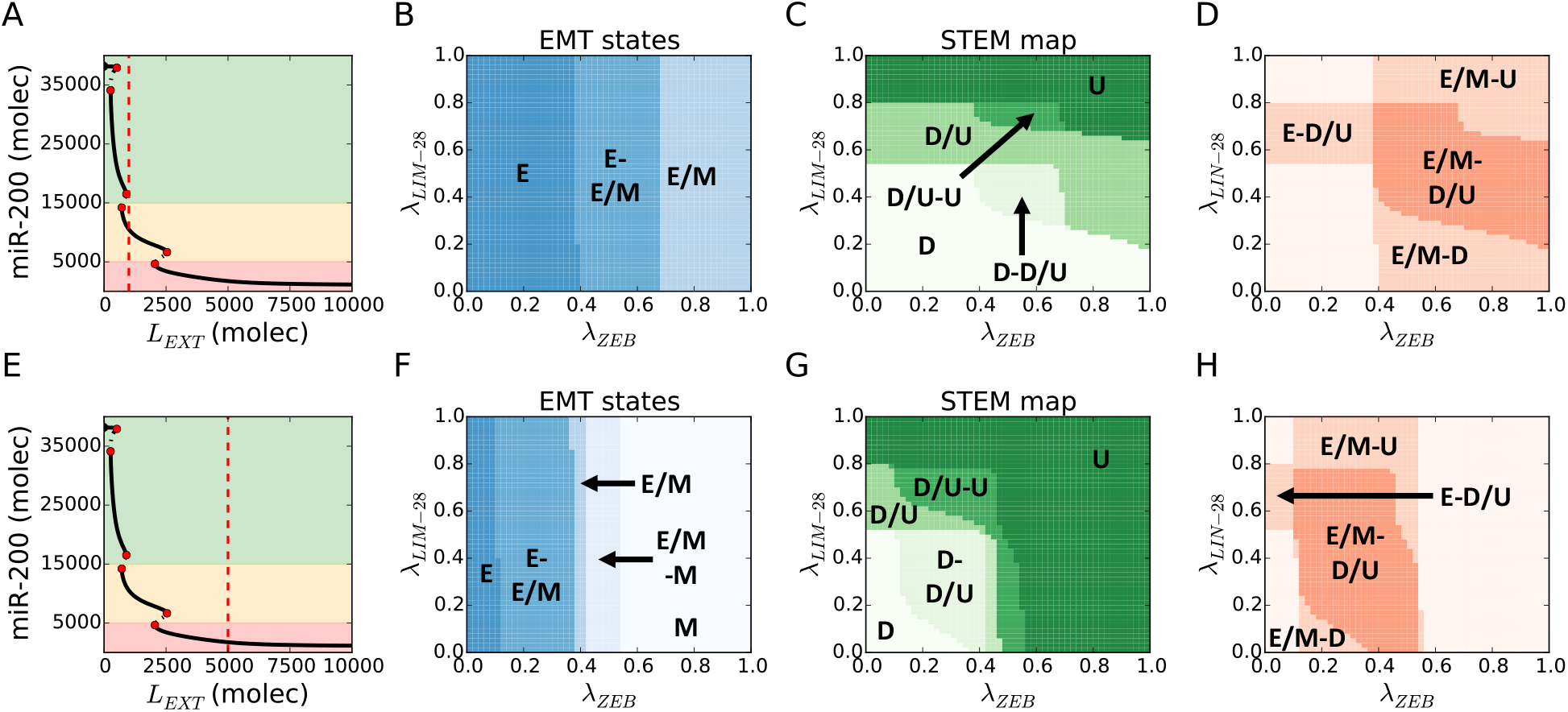
Hybrid E/M and D/U (stem-like) phenotypes overlap over a large variation of EMT-STEM circuit coupling strength. **(A)** For *L_EXT_* = 10^3^ molecules and no EMT-STEM coupling *λ*_*LIM*–28_ = *λ_ZEB_* = 1 (first considered condition), the cell expresses a hybrid E/M state (see the intersection of the red dotted line with the diagram’s continuous lines). **(B)** EMT state diagram over variation of fold-changes for let-7 inhibition on ZEB (*λ_ZEB_*, x-axis) and micro-RNA-200 inhibition on LIN-28 (*λ*_*LIM*–28_, y-axis) and for *L_EXT_* = 10^3^ (panel A). The cell phenotype is described by the state of the EMT module. **(C)** STEM phenotypic characterization diagram for the conditions of panel A. The cell phenotype is described by the state of the STEM module (Down: D, Stem: STEM, Up: U). **(D)** Superimposition of the stability regions of hybrid E/M state (panel B) and D/U state (panel C) for the conditions of panel A. Labels highlight the regions where the hybrid E/M and/or stem-like D/U are observed. This diagram does not label every combination of EMT and STEM phenotypes but only highlight regions where either hybrid E/M or stem-like D/U (or both) are expressed. **(E)** For *L_EXT_* = 5 10^3^ molecules and no EMT-STEM coupling *λ*_*LIM*–28_ = *λ_ZEB_* = 1 (second considered condition), the cell expresses a monostable mesenchymal phenotype. **(F)** EMT state diagram for *L_EXT_* = 5 10^3^ (panel E). **(G)** STEM phenotypic characterization diagram for the conditions of panel E. **(H)** Superimposition of the stability regions of hybrid E/M state (panel F) and D/U state (panel G) for the conditions of panel E.

We repeat the abovementioned analysis, when the cell is exposed to a high level of external Notch ligands, and therefore in a mesenchymal state initially (due to Notch-induced EMT) (Fig. 3E). In this case, all three EMT states (E, E/M, M) and STEM phenotypes (DOWN, D/U, UP) are observed upon variation of the coupling parameters (*λ*_*LIM*–28_, *λ_ZEB_*) (Fig. 3F-G). Similar to the first case, the regions of the hybrid E/M state and the D/U phenotype largely overlap (darker region in Fig. 3H), hence confirming the correlation reported in the Notch-driven EMT. Further, very strong repression of ZEB by let-7 facilitates the existence of epithelial stem-like cells (i.e. {E - D/U}), while very weak repression of ZEB by let-7 enables the overlap of mesenchymal state with stem-like behavior (i.e. {M-D/U}) (*λ_ZEB_* close to 0 and close to 1 in Fig. 3F respectively).

Overall, these results consistently show a strong correlation between hybrid E/M and D/U stem-like traits across large variation of the coupling strength between EMT and STEM circuits. Further, other factors such as the activation status of Notch signaling can shift the ‘stemness window’ across the EMT axis, resulting in epithelial (E-D/U) or mesenchymal (M-D/U) stem cells. Thus, while the ‘stemness window’ may be likely to lie mid-way on the EMT axis, context-specific differences may shift it towards either end (i.e. E or M) of the EMT axis.

### Tuning NF-kB, EMT induction, and Notch activation levels reveal different subpopulations of Cancer Stem Cells (CSCs)

Crosstalk with intra and/or extra-cellular signaling pathways can modulate the NOTCH, EMT and STEM modules, thus altering their expected outcomes. To incorporate this aspect, we consider three signals that can serve as inputs for the NOTCH-EMT-STEM circuit: (i) external Notch ligands (Delta and Jagged) *L_EXT_* activating intra-cellular Notch signaling; (ii) a direct EMT-inducer *I_EXT_* that may stabilize or overexpress SNAIL, such as Wnt or TGF-beta; and (iii) NF-kB signaling activating LIN-28 and let-7[13].

To evaluate the joint effect of *L_EXT_*, *I_EXT_* and NF-kB on the region enabling the (high notch, high jagged)-hybrid E/M-stem-like phenotype – or the coupled ‘E/M - D/U - S/R’ state, we set up three phenotype characterization diagrams where two parameters are varied while the third is held constant, and inspect the parameter variation range over which the coupled ‘S/R – E/M – D/U’ phenotype exists. For this set of simulations, the connection between EMT and STEM modules is intermediate (*λ*_*LIM*–28_ = *λ_ZEB_* = 0.5, the central point in the diagrams of Fig. 3).

First, simulating the coupled modules at a fixed value of NF-kB, we found that the ‘S/R – E/M – D/U’ phenotype exists for intermediate levels of exposure to Notch ligands (*L_EXT_*) while exposure to EMT induction signals (*I_EXT_*) narrows the stability range of the ‘S/R – E/M – D/U’ phenotype (Fig S2). Further, fixing the value of EMT inducer *I_EXT_* shows that a strong down-regulation or overexpression of NF-kB restricts the stability of the ‘S/R – E/M – D/U’ phenotype as well (Fig. S3), i.e. the stability range of ‘E/M – D/U – S/R’ phenotype is maximized at intermediate values of NF-kB.

Next, simulating the coupled modules at fixed values of *L_EXT_*, we observed that a strong EMT-induction (*I_EXT_*) pushes the cell out of the (high Notch, high Jagged) S/R region and into a mesenchymal (M) state, while NF-kB overexpression could rescue the existence of S/R and E/M phenotypes (Fig. 4A-B). Further, LIN-28 is upregulated in presence of high NF-kB, leading to an UP phenotype (Fig. 4C). Overall, high levels of EMT-induction and NF-kB push the cell out of the coupled ‘E/M – D/U – S/R’ phenotype (Fig. 4D). Specifically, a strong EMT induction can result in mesenchymal stem cells (M-D/U at high *I_EXT_*, low NF-kB), while NF-kB overexpression can generate hybrid E/M that are not stem-like (E/M-U at low *I_EXT_*, high NF-kB). Finally, we observed non-stem mesenchymal cells when both signaling channels – *I_EXT_* ad NF-kB – are active (M-U at high *I_EXT_*, high NF-kB).

**Figure 4.**
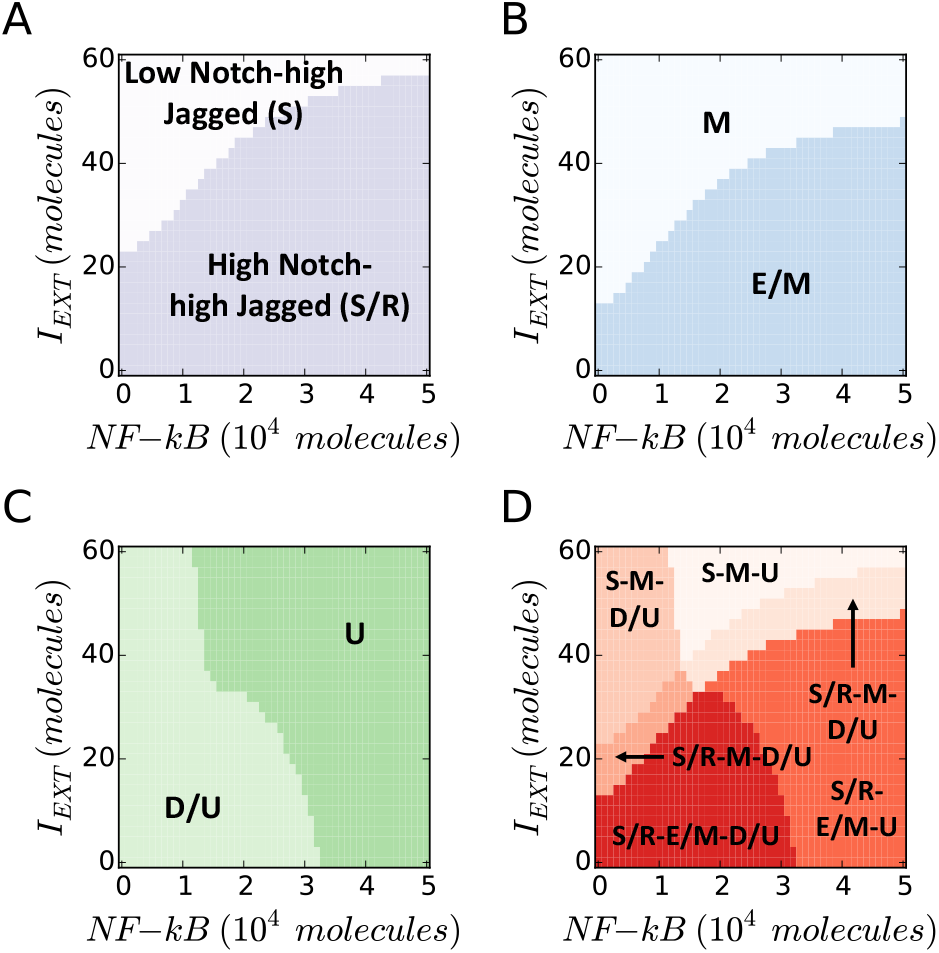
EMT Induction and NF-kB overexpression push cancer cells out of the “S/R-E/M-D/U window”. **(A)** Phenotypic characterization diagram of the Notch phenotype in presence of variable NF-kB (x-axis) and EMT-Inducer *I_EXT_* (y-axis). A high *I_EXT_* pushes the cell out of the (high Notch, high Jagged) S/R phenotype, while NF-kB increases the *I_EXT_* threshold needed to exit the S/R phenotype. **(B)** State diagram of the EMT state. The cell transitions from hybrid (E/M) to mesenchymal (M) when I_EXT_ is increased, while NF-kB increases the *I*_*EXT*_ threshold required for the transition. **(C)** Phenotypic characterization diagram of the Stem phenotype. The cell switches from D/U – or STEM – to UP when NF-kB is increased, while *I_EXT_* decreases the NF-kB threshold required for the transition. **(D)** Overlap of the three maps highlights the S/R-E/M-D/U window. A large *I_EXT_* and/or overexpression of NF-kB pushes the cell out of the window. In this simulation, the cell phenotype is measured upon full equilibration. The initial condition is always within the S/R-E/M-D/U window. *L_EXT_* is constant at 2000 molecules and the EMT-STEM coupling is intermediate (*λ*_*LIM*–28_ = *λ_ZEB_* = 0.5).

All possible combinations of stem and non-stem cells undergoing partial or complete EMT were observed, thereby showing that the coupling among EMT, Notch, and STEM modules can give rise to different subpopulations of cancer stem cells (CSCs) with a spectrum of epithelial-mesenchymal phenotypes.

There is every reason to suspect that the precise set of phenotypes seen in a given situation depends on the cancer type, it is probably very specific from patient to patient, and perhaps even depend on the specific position within a given tumor[36,37].

### Analysis of gene expression profiles reveals epithelial, hybrid E/M and mesenchymal Cancer Stem Cells

To validate our prediction that context-specific interactions can move the ‘stemness window’ toward the epithelial or mesenchymal end of the ‘EMT axis’, we analyze the gene expression data of CSCs from different cancer subtypes. We previously devised an inferential model which predicts the positioning of a given gene expression profile along the ‘EMT axis’, or ‘EMT score’, using the expression level of several key EMT regulators as predictors[38]. Among others, the EMT metric score considers canonical epithelial and mesenchymal markers such as E-cadherin and Vimentin as well as ‘phenotypic stability factors’ (PSFs) which help in stabilizing a hybrid E/M phenotype such as GRHL2 and OVOL. These scores are on a scale of 0 (fully epithelial) to 2 (fully mesenchymal).

CSCs isolated from various non-small cell lung cancer cells – A549 and NCI-H2170 – were identified as either hybrid E/M or mesenchymal. Furthermore, CSCs from breast cancer (MCF7), thyroid cancer, and glioblastoma were classified as hybrid E/M, epithelial, and mesenchymal respectively, highlighting the significant heterogeneity in EMT status of CSCs isolated from varying cancer types (Fig 5). This heterogeneity is also reflected via analyzing the data from spheres formed by colorectal cells (H29), glioma cells, and by heterogeneous HMLER cells. Finally, the side-population in pancreatic adenocarcinoma that express many CSC genes was predominantly hybrid E/M (Fig 5). Put together, these strongly suggest that while stem-like properties are most likely to be associated with a hybrid E/M phenotype, in accordance with our modeling prediction, the possibility of epithelial or mesenchymal subsets of CSCs is not ruled out. It is worth pointing out that since the datasets used here for CSCs are from a cell population instead of single-cell, a hybrid E/M phenotype prediction may correspond to *bona fide* hybrid E/M cells and/or a mixture of E and M cells.

**Figure 5.**
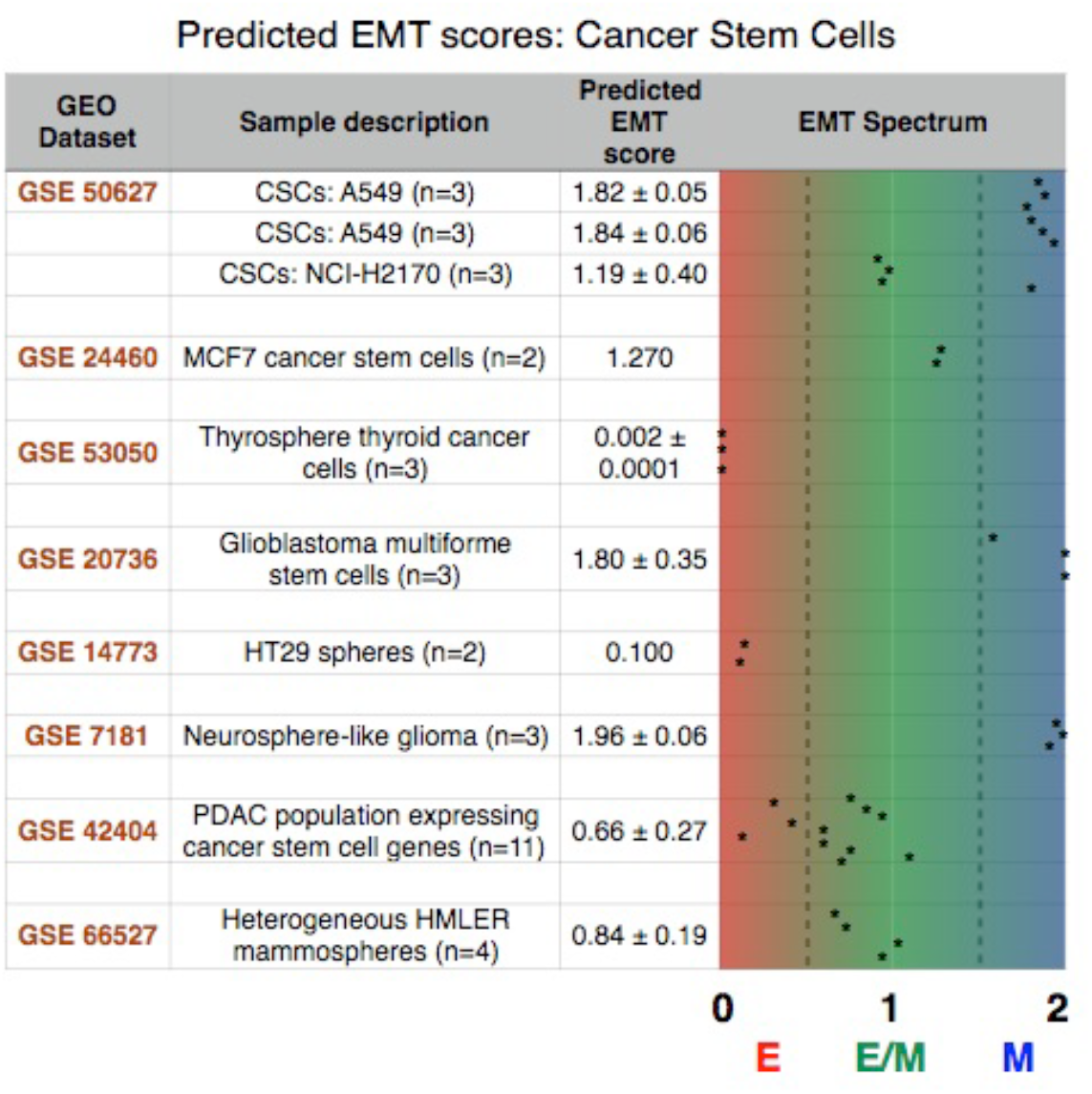
The predicted EMT score for several stem cancer subtypes shows a heterogeneous distribution across the EMT axis. Each dataset is identified by its GEO number. The number of each sample in a given dataset along with a brief explanation is provided in sample description. Mean and, when applicable, standard deviation for relevant samples in each dataset are reported in the predicted EMT score category, and individual samples are graphed on the EMT spectrum to illustrate sample heterogeneity.

### Metformin restricts the existence of coupled E/M- D/U- S/R state

To further test the ability of our computational model in recapitulating experimental observations, we model the effect of metformin in decreasing tumor aggressiveness and targeting CSCs. Metformin, the most widely used anti-diabetic drug, is recently receiving attention as an anticancer drug in the context of several cancer types, including prostate, breast, lung, and pancreatic cancer [39,40]. Among other observed effects, it selectively targets CSCs growth [41,42] via antagonizing LIN28 [43], decreases EMT and cell invasiveness in melanoma[44], inhibits TGF-beta induced EMT in cervical carcinoma cells [28], and reduces metastases in mice models [42]. The assessment of the effect of Metformin can in the future be applied other drug treatments that couple to the basic elements of our circuit.

We integrate metformin into our mathematical model as an inhibitor of SNAIL and LIN-28 to consider the EMT-halting and CSC targeting effects, respectively [28,43] (see Methods), and re-compute the (*I_EXT_*, NF-kB) phase diagram shown in Fig. 4.

We find that in presence of metformin, the (high Notch, high Jagged), i.e. hybrid S/R phenotype is accessible only under a strong EMT induction (Fig. 6A). This result is consistent with the decrease in Notch1 levels in metformin-treated pancreatic cancer cells [27]. Metformin also acts as an EMT brake by inhibiting SNAIL, therefore demanding a stronger EMT push to achieve a hybrid E/M state (Fig. 6B).This finding agrees well with its halting effect on TGF-beta induced EMT[28]. At low levels of EMT inducing signals, the cell is epithelial (E) and Notch signaling is therefore strongly inhibited via post-translational regulation of Notch signaling components by microRNAs miR-34 and miR-200. Thus, the effect of metformin on Notch levels can be explained through the coupling between Notch and EMT modules. Furthermore, under the effect of metformin, the cell attains a DOWN/UP, i.e. stem-like phenotype only at very strong EMT induction and NF-kB overexpression (Fig. 6C). In other words, our model predicts that metformin treatment pushes a large fraction of cells outside the ‘stemness window’, a prediction which is consistent with its effect in targeting CSC via LIN-28. Combining these results shows that metformin severely restricts the coupled ‘E/M- D/U- S/R’ phenotype; this state is rescued only under a very strong EMT induction and NF-kB overexpression (Fig. 6D).

**Figure 6.**
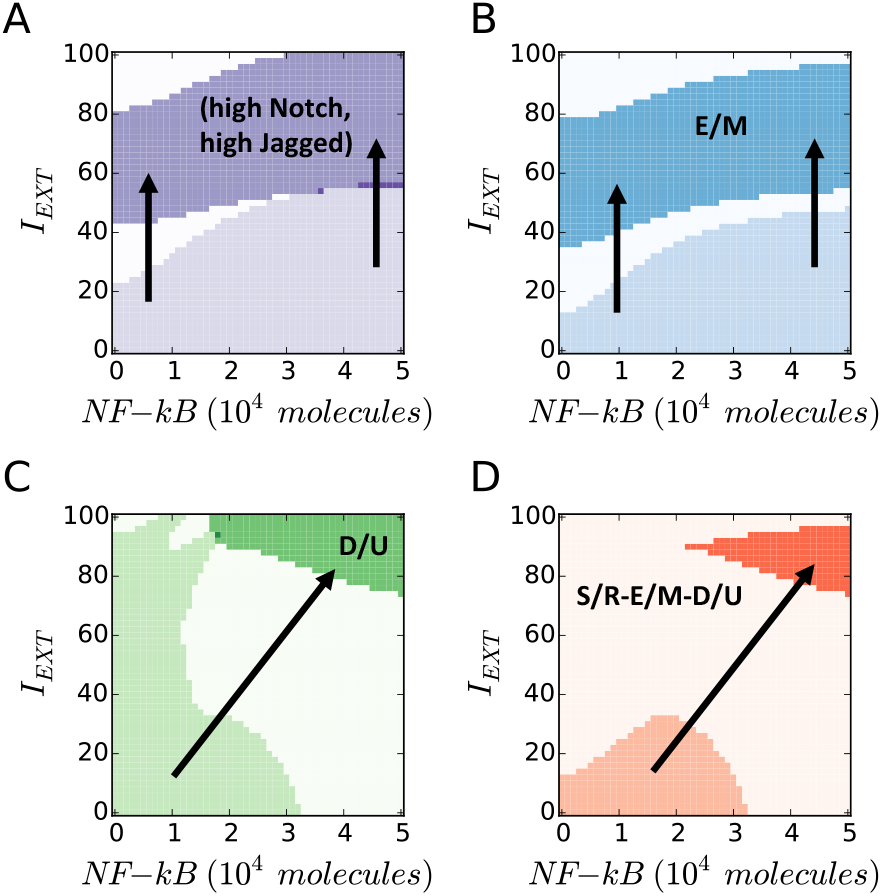
Metformin restricts the existence of coupled E/M- D/U- S/R state. **(A)** The (high Notch, high Jagged) region is shifted to higher levels of external EMT-inducer *I_EXT_* in presence of metformin (dark purple) compared to the control (i.e. no metformin) case (light purple). **(B)** The hybrid E/M EMT state is shifted to higher levels of external EMT-inducer *I_EXT_* in presence of metformin (dark blue) compared to the control case (light blue). **(C)** The D/U STEM phenotype is pushed to (high EMT-inducer, high NF-kB) in presence of metformin compared to the control case (dark green vs light green). **(D)** Metformin enables the existence of a coupled ‘E/M – D/U- S/R’ state only at (high EMT-inducer, high NF-kB) levels (dark orange), as compared to the control case where the ‘E/M- D/U- S/R’ state can exist at (low EMT inducer, low NF-kB) levels (light orange). For this simulation, *L_EXT_* = 2000 molecules and the EMT-STEM coupling is intermediate (*λ*_*LIM*–28_ = *λ_ZEB_* = 0.5).

## Discussion

We introduced a mechanism-based mathematical framework to investigate the interplay between epithelial-mesenchymal transition (EMT), cancer stem cells (CSCs) and Notch signaling - three key axes contributing to cancer metastases and therapeutic resistance [4,18,45].

Our model suggests a strong correlation between a hybrid E/M phenotype with CSC properties and augmented Notch-Jagged signaling. Such finding resonates well with the increasing evidence indicating that the higher plasticity and multi-lineage differentiation potential are more likely to be found midway *en route* to EMT – and not towards the mesenchymal end [46–48]. Furthermore, the predicted hybrid E/M stem-like state associates with reinforced Notch-Jagged signaling that (a) promotes resistance against chemo- and radio-therapy, (b) facilitates colonization by promoting cell-cell communication between the cancer cells and the cells of the organ where they extravasate, as seen during bone metastases of breast cancer[49], and (c) coordinates spatial co-localization of hybrid E/M cells in the tumor tissue, hence facilitating the formation of clusters of hybrid E/M cells [4]. Furthermore, Jagged1 - both in its transmembrane form (present on the cell surface) and soluble form (secreted by stromal endothelial cells) – appear to be a potent inducer of Notch signaling in maintaining and expanding the CSC population [20,21]. Consequently, Jagged1 levels are overexpressed in CSCs as compared to non-CSCs [50,51], and are associated with poor survival and recurrence [18]. Overall, these findings support the increasingly accepted notion that a hybrid E/M state – and not necessarily a completely mesenchymal M state – should be considered as a hallmark of cancer aggressiveness [52,53].

We further predict that while the ‘stemness window’ is most likely to lie midway along the ‘EMT axis’, various external signals and/or varying strengths of interactions among the Notch, EMT and STEM modules may shift the window along the axis. This prediction is supported by our analysis of gene expression profiles of CSCs belonging to different cancer subtypes, and report the presence of epithelial, hybrid E/M and mesenchymal CSCs. Recent experimental studies have identified such heterogeneity in EMT status of different CSCs across cancer subtypes [10,12,36,54–57], supporting the idea about a dynamic ‘stemness window’[13] along the ‘EMT axis’. Moreover, a recent clinical study identified different subsets of CTCs - those that expressed EMT markers but not stemness ones, those that expressed stemness markers but not EMT, those that expressed both EMT and stemness markers, and those that expressed neither. The relative frequencies of these subsets changed upon neoadjuvant chemotherapy [57]. Put together, these observations display the importance of context-specific factors in modulating the phenotype of a cancer cell on the EMT and stemness axes [58,59], including therapy-induced adaptive phenotypic transitions[60]. For instance, micro environmental factors such as inflammation and hypoxia can activate EMT and therefore reinforce a CSC phenotype[46]. Further, Notch ligands from neighboring cells (juxtacrine) or in soluble form (paracrine) can enhance the activation of Notch signaling, thereby boosting drug resistance, activating EMT and promoting CSCs [61,62]. Indeed, human breast stem cells co-expressing various epithelial and mesenchymal markers, as identified by single-cell RNA-seq, exhibited enriched Notch signaling. Consistently, Notch3 was found to be overexpressed in highly aggressive triple negative breast cancer samples and correlated with poor patient survival [37].

Previous mathematical models and experimental studies have identified multiple phenotypic stability factors (PSF) that can stabilize a hybrid E/M phenotype such as OVOL1/2[63–65], GRHL2[66], ∆NP63α[67], and NUMB/NUMBL [68] and facilitate collective cell migration. GRLH2 and OVOL1/2 directly target the miR-200-ZEB axis regulating EMT[63,66], while NUMB/NUMBL modulates EMT via Notch-Jagged signaling[68]. Higher levels of these PSFs can also correlate with poor patient outcome [69]. Thus, future studies should investigate the effects of these and other PSFs in regulating cancer cell aggressiveness *in vitro* and *in vivo* to further elevate our understanding of the connection between EMT and stemness, and might potentially provide novel therapeutic targets to break the clusters of circulating tumor cells (CTCs) driving metastases.

Lastly, the model recapitulates the effect of metformin, a drug capable of inhibiting EMT and selectively targeting CSCs[28,41]. Its effect can be interpreted in terms of restricting the existence of a coupled ‘S/R - E/M - D/U’ state. In particular, reversing a partial EMT naturally restricts the Notch-Jagged signaling axis [27], while overexpressing NF-kB recovers stem-like traits via activation of LIN-28 inhibited by metformin[29].

Therefore, our model provides a mechanism-based explanation for otherwise uncorrelated biological observations among Notch, EMT and stem-like traits, and offers a predictive platform towards gaining an integrative, functional, mechanism-based understanding of cancer metastases.

## Methods

### Notch module

The level of Notch receptor (N), Delta (D) Jagged (J) and NICD (I) in the cell are modeled via a system of ordinary differential equations according to the model of Boareto *et al*.[25,70]:

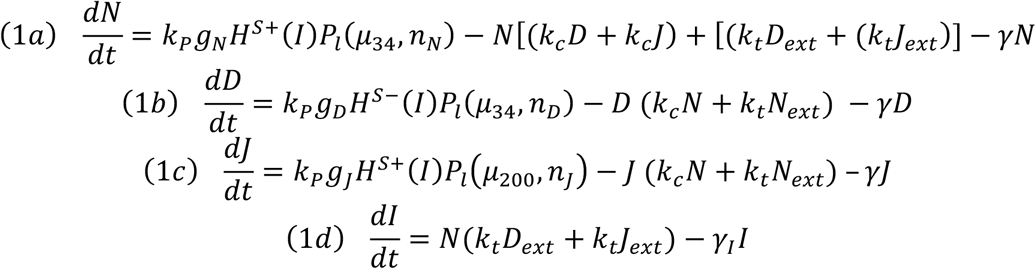

with transcription rates *g_N_*, *g_D_*, *g_J_* for Notch, Delta and Jagged mRNA (not explicitly present in the model) and translation rate *k_p_*. Notch, Delta and Jagged degrade at rate *γ*, while NICD has a faster rate *γ_I_*. The functions *H*^S+^(*I*)/*H*^S−^ (*I*) indicate positive/negative transcriptional regulation of NICD activating Notch, Jagged and inhibiting Delta (see Supplementary Information section “Mathematical modeling of transcriptional/translational interactions” for details). The function *P_l_*(*μ*, *n*) models the post-translational inhibition exerted by micro-RNA *μ* (miR-34 or miR-200) binding on the *n* binding domains on Notch, Delta or Jagged. These terms therefore represent the connection between EMT and Notch modules (see Supplementary Information section “Mathematical modeling of post-translational interactions” for details). *k_c_* and *k_t_* are the receptor-ligand binding constants for cis-interaction (receptor and ligand from same cell leading to complex degradation) and trans-interaction (native receptor binding with external ligand leading to NICD release). *N_ext_*, *D_ext_* and *J_ext_* are the amount of external Notch, Delta and Jagged at cell surface available to bind with the cell’s receptors and ligands.

The phenotypes expressed by the Notch module are based on the levels of Notch receptor and Jagged ligand. We introduce thresholds for the Notch receptor (N~13000 receptor molecules) and the Jagged ligand (J~350 ligand molecules) such that the (high Notch, high Jagged) hybrid Sender-Receiver (S/R) phenotype satisfies (Notch > N, Jagged > J). If (Notch < N, Jagged > J) the cell is a (low Notch, high Jagged) Sender (S) cell, while if (Notch > N, Jagged < J) the cell is a (high Notch, low Jagged) Receiver (R) cell. In principle, an inactive (Notch < N, Jagged < J) phenotype should also be considered but is never observed for the chosen values of N, J. Previous modeling on the Notch-Delta-Jagged system by Boareto *et al*.[25,70] defined the Notch phenotypes based on the branches of a bifurcation diagram, similar to the current definition of the EMT states. Therefore, our model does not classify the Notch phenotypes in the same way but maintains the same terminology (Sender, Receiver, hybrid Sender/Receiver).

### EMT module

The interactions between miR-34 (*μ*_34_), miR-200 (*μ*_200_), ZEB (Z) and SNAIL (S) depicted in Fig. 1B are modeled via a system of ordinary differential equations according to Lu *et al*.[71]:

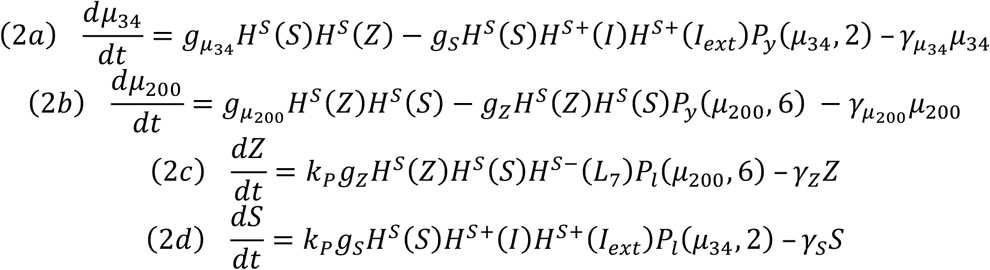

with basal transcription rates (*g*_*μ*_200__, *g*_*μ*_34__, *g*_*Z*_, *g*_*S*_), translation rate *k_P_* and degradation rates (*γ_μ_200__*, *γ_μ_34__*, *γ_Z_*, *γ_S_*). Similar to the Notch module, the functions *H^S+^(I)/H^S−^(I)* model transcription/translational interactions of ZEB and SNAIL while the function *P_l_(μ, n)* represents post-translational inhibition of SNAIL and ZEB by miR-34 and miR-200, respectively. The corresponding loss of miR-34 and miR-200 due to micro-RNA-protein complex degradation is modeled via the associate function *P_y_*(*μ*, *n*) (see Supplementary Information section “Mathematical modeling of post-translational interactions” for details). The term *H^S+^*(*I*) represents transcriptional activation of SNAIL by NICD, and therefore connects the Notch and EMT modules. Additionally, the term *H^S^*^−^(*L*_7_) models the inhibition of ZEB by Let-7, thereby connecting Stem and EMT modules. The effect of an external EMT inducer *I_ext_* is considered via the shifted Hill function *H^S^*^+^(*I_ext_*) activating SNAIL.

The EMT states (epithelial, hybrid E/M, mesenchymal) are defined based on the levels of micro-RNA miR-200 (epithelial: miR-200 > 15000 molecules; hybrid E/M: 5000 molecules < miR-200 < 15000 molecules; mesenchymal: miR-200 < 5000 molecules). This definition reflects the mathematical solutions, or branches, of the bifurcation diagram of Fig. 2A, and was already used in the original work by Lu *et al*.[71] that introduced the EMT module.

### Stem module

The stem circuit of Fig. 1C including LIM-28 (*L*_28_) and Let-7 (*L*_7_) is described by the model of Jolly et al.[13]:

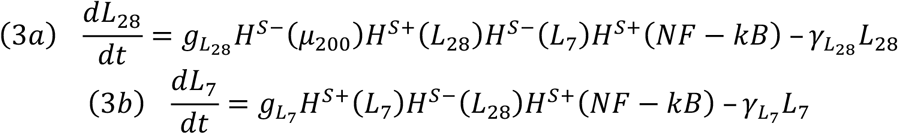

Where (*g_L_28__*) and (*g_L_7__*) are production rates and (*γ_L_28__*) and (*γ_L_7__*) are degradation rates. In the genetic circuit, LIM-28 self-activates (*H^S+^*(*L*_28_)) and inhibits Let-7 (*H^S–^*(*L*_7_)). Similarly, Let-7 activates itself (*H^S+^*(*L*_7_)) and represses the expression of LIM-28 (*H^S–^*(*L*_28_)) resulting in the double negative switch. The stem module is connected to the EMT module via the term *H^S–^*(*μ*_200_) describing inhibition of LIM-28 by miR-200. Finally, *H^S^*^+^(*NF* − *kB*) represents the effect of NF-kB signaling activating Lim-28 and Let-7.

The stem interval (as shown in Fig. 2B) is defined as follow. We considered the extremal values assumed by LIN-28 (the minimum m at *L_ext_* = 0 molecules and the maximal M at *L_ext_* = 10000 molecules) and defined the window as [ m+0.25(M-m), m+0.65(M-m)] [13,15]. Based on the diagram of Fig. 2B, *m*~56000 molecules, *M*~110000 molecules. Therefore, the cell is DOWN for LIN-28 < m, DOWN/UP for m < LIN-28 < M, and UP for LIN-28 > M. The motivation for this classification is that an intermediate expression level of OCT4, a direct target of LIN-28, has been associated with stem-like traits[13].

### Numerical calculation details

We developed all source code in Python and used the numerical library PyDSTool[72] to compute the bifurcation diagrams. All plotting was performed using the Python numerical library Matplotlib[73].

### EMT score quantification

The EMT Metric previously described[38] was applied to various Gene Expression Omnibus (GEO) datasets containing CSCs. A collection of EMT-relevant predictor transcripts as well as a cross-platform normalizer transcripts was extracted for each dataset and used to probabilistically categorize samples into an element of {E, E/M, M}. To each sample i there corresponds an ordered triple S_i_=(P_E_, P_E/M_, P_M_) that characterizes the probability of group membership. Categorization was assigned based on the maximal value of this ordered triple. S_i_ was then projected onto [0,2] by use of the EMT metric. The metric places epithelial (resp. mesenchymal) samples close to 0 (resp. 2) while maximally hybrid E/M samples are assigned values close to 1.

